# Design and characterization of SAKe, a new building block for protein self-assembly

**DOI:** 10.64898/2026.03.16.710736

**Authors:** Andreu Mor Maldonado, Staf M.L. Wouters, Hiroki Noguchi, David E. Clarke, Gangamallaiah Velpula, Steven De Feyter, Arnout R. D. Voet

## Abstract

The nanofabrication of functional protein-based surfaces is challenging due to the chemical complexity of proteins and their unpredictable behavior at the solid-liquid interface. Many proteins of interest –such as antibodies or large enzymatic complexes – lack strong and dynamic protein-protein and protein-surface interactions necessary to drive self-assembly of stable arrays with high surface coverage. Additionally, adsorption-induced conformational changes at the solid-liquid interface could lead to a loss of activity and increase the risk of undesirable interfacial processes. Here we introduce SAKe, a kelch-like designer protein, as a versatile platform to address these challenges. Ancestral sequence reconstruction led to high thermal stability, and the high symmetry allowed modularity of the protein’s core. Rational engineering of the bottom side allowed SAKe to form large (up to 5 micrometers in length), well-defined and pH-dependent two-dimensional assemblies while maintaining structural integrity, which is key for further development of functional materials. SAKe self-assembly was investigated through in-liquid atomic force microscopy on muscovite mica. High resolution imaging confirmed the integrity of the SAKe protein upon adsorption on the solid-liquid interface. These results showcase the SAKe protein as a platform for the further engineering of functional protein-based two-dimensional materials.

## INTRODUCTION

Nanotechnology aims to achieve precise control over functional nanomaterials at the atomic scale.^1^ Biological systems, particularly proteins, offer a promising starting point due to their chemical variability. Common strategies to functionalize relevant surfaces such as muscovite mica and graphene are direct adsorption or covalent linkage to reactive moieties such as thiols, amines or carboxylic groups.^2–6^ For example, the group of Duncan covalently immobilized plasma fibronectin onto polystyrene via thiol or amine groups. While this resulted in improved chemical stability, it did not produce a well-organized protein layer, illustrated by high heterogeneity in the z-plane.^5^ To improve spatial control over the localization of reactive groups on the surface, a scaffolding protein layer can be used. A well-studied model system of this kind is the S-layer protein (SLP).^6–8^ S-layer proteins readily form two-dimensional arrays with tunable periodicity. These proteins have been successfully used for the functionalization of surfaces with both biomolecules and inorganic materials such as metal nanoparticles.^9^

While covalent linkage results in great stability of the proteins at the interface, it relies on highly reactive groups that are broadly distributed on the protein surface. Such little control over the interaction between the protein and the interface leads to unpredictable and heterogeneous protein orientation at the solid-liquid interface, which relates to low surface coverage.^5,10,11^ Moreover, covalent attachment of proteins at interfaces directly impacts a key determinant of crystallinity at the solid-liquid interface. Covalent bonds prevent molecular diffusion and molecular rearrangement on the surface, impeding proteins from adopting the correct orientation with respect to nucleation sites, thereby hindering the growth of defect-free crystalline arrays.^12^ The need of such dynamic behavior is clearly seen in the formation of SLP arrays. The group of Magalí described such dynamic behavior imaging three different phases for the formation of a crystalline layer of SLP on muscovite mica. They reported that initially proteins are randomly adsorbed on the surface, then the nucleation sites start growing reaching a final stage where proteins suffer a conformational change to stabilize the assemblies. These processes required up to one hour to reach the final stable assembly.^13^

Physisorption of the proteins onto the surface, on the other hand, often leads to protein denaturation as well as heterogeneity of possible protein orientations at the solid-liquid interface; especially when using hydrophobic surfaces.^14,15^ The group of Craighead directly adsorbed ConA (a lectin protein) on graphene. This immobilization resulted in the loss of the binding capacity to oligosaccharides.^10^ Thus, showcasing how arbitrary orientation of the proteins at the interface limits accessibility of their functional sites. Additionally, inefficient surface coverage and protein unfolding leave open areas for non-specific interactions which may decrease the sensitivity of biosensing, or lead to undesired by-products in catalysis.^6,14–16^

Rational protein engineering can increase surface coverage and homogenize protein orientation. For example, the groups of Tezcan and Zhan achieved highly symmetrical protein assemblies by introducing disulfide bonds or metal coordination sites to L-Rhamnulose-1-phosphate aldolase and tobacco virus coat protein respectively.^17,18^ Ju Song and coworkers used non-canonical amino acids to drive protein self-assembly. 2D crystals were assembled in solution by placing bipyridyl-alanine groups at geometrically interesting positions in a D3 homohexamer.^19^ Lastly, efficient protein surface assembly can be achieved by computational redesign of either protein-protein interfaces or protein-surface interfaces. This was demonstrated by the design of an α-helical repeat protein capable of highly efficient self-assembling on mica by the Baker group.^20,21^

A common trait in rationally designed self-assembling systems is the use of protein symmetry. Near-perfect symmetry is commonly found in proteins and especially in oligomers.^22–24^ It is hypothesized that symmetry facilitates protein folding and has a strong correlation with protein function. The geometry of protein complexes can be controlled by tuning the symmetry of the building blocks.^22,25–27^ Symmetric placement of protein contact points largely facilitates the interaction between molecules as well as reduces the possible conformations that proteins can adopt within the complex.^25^

While protein engineering has made significant advances in the nanofabrication of two-dimensional materials, current approaches still depend on complex protein modifications or on the use of intermediate molecules and/or scaffolding proteins to achieve high surface coverage. Despite intensive effort, protein scaffolds capable of forming densely packed two-dimensional arrays combined with the ability to implement a desired function remain underexplored.

Our group previously designed and engineered a fully symmetric beta propeller, Pizza.^28^ This scaffold was further engineered for polyoxometalate coordination,^29^ metal-free catalysis^30^ as well as to scaffold the crystallization of salt nanocrystals.^31^ Despite these varied applications, the application potential of Pizza was limited given the lack of a large flexible protein surface. Following the same design principles used in the engineering of Pizza,^28^ a new pseudo-symmetric protein scaffold was used as a starting point, this time targeting proteins mediating protein-protein interactions as their natural function.

Here, we report on the computational design and biophysical characterization of a modular protein building block (SAKe) inspired by the kelch protein family. SAKe proteins were designed as candidate proteins for achieving this dual requirement of self-assembly and function. These proteins not only show great self-assembling properties both in solution and on surface but also can tolerate extensive mutations to further develop them into functional building blocks.

## RESULTS AND DISCUSSION

### Design and characterization of the SAKe Protein Scaffold

The Kelch folding domain was used as a starting point for the design of a stable symmetric scaffold. Kelch repeat proteins are β-propeller proteins composed of 6 nearly identical tandem sequence repeats that each fold into four-stranded anti-parallel sheets (referred to as blades) around a central cavity.^32,33^ While the core of these blades is well conserved, the top-side loops vary both in sequence and length **(Figure 1)**, influencing the protein’s function. The majority of natural Kelch domains modulate protein-protein interactions (PPIs), especially in humans as an adaptor protein of the CUL3 complex.^34^ However, the loops which mediate the protein interactions as observed in the CUL complexes on some Kelch domains are also found to harbor an enzymatic site, demonstrating the versatility of this protein fold.^32,33,35^

**Figure 1:**
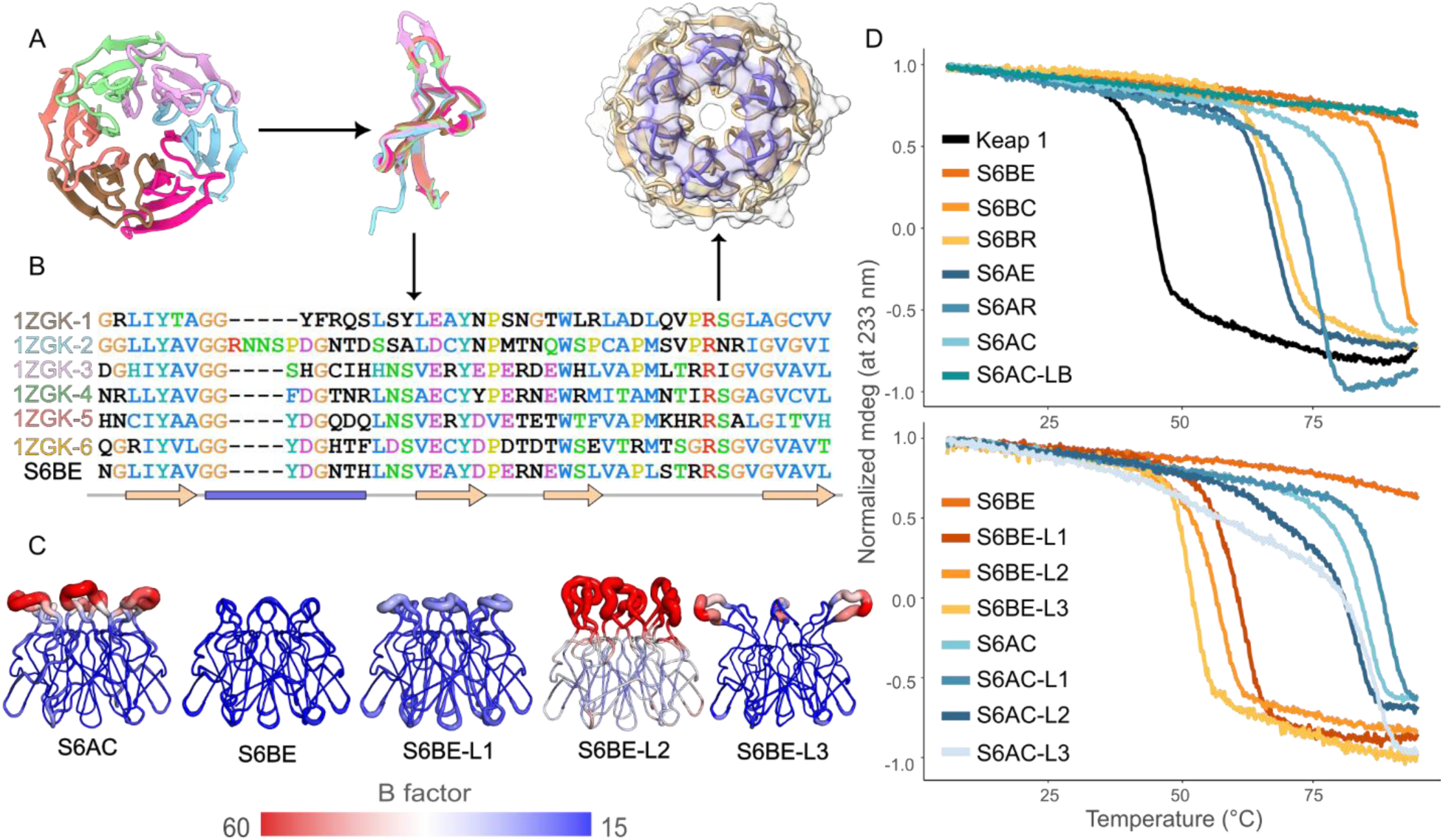
SAKe design strategy. The six blades of Keap 1 kelch domain (1ZGK) were isolated **(A)** and used to construct an MSA **(B)** in order to create a fully symmetrical kelch protein using the RE3Volutionary method. The B-factors of the different loop lengths indicate the high flexibility of the loops as the length increases **(C)**. The high stability of the SAKe protein is reflected by very high melting temperatures. As the loop length increases, the flexibility of the top loops increases, what notably reduces the thermal stability of the proteins **(D)**.

The Kelch domain of the Keap1 protein was selected as the starting point for proteins design as RADAR analysis^36^ revealed it to consist of the most conserved repeats of all crystalized Kelch domains exhibiting a highly symmetrical tertiary structure. Proteins with a high degree of symmetry have not only been shown to have superior stability to their non-symmetric counterparts, but it has shown also to play a key role in molecular self-assembly.^25,37,38^ Designing symmetrical building blocks is important as the number of available contact points to drive and stabilize the complex increases equally around the molecule. This is key as the structure of the final oligomer is often determined by the initial seeding of the monomers.^37,39,40^

Next, an ensemble of SAKe proteins was generated from the human Keap1 β-propeller using the RE_3_Volutionary design procedure **(Figure 1).**^28^ Because the Keap1 template is composed of blades carrying loops of three different lengths, Rosetta symmetry docking was used to generate three different C_6_ symmetry starting backbones.^41^ All models retained a velcro closure type identical to the parent Keap1 protein^42^ **(Figure S1).**

Initially, the backbones from the 2^nd^ and 6^th^ blades of the Keap1 kelch domain were chosen to generate the type A and type B SAKe backbones, respectively. These two blades were chosen as initial design backbones since they accommodate the most common loop length (six amino acids) observed in human propellers. This way, the two most common loop lengths of the Keap1 kelch repeats could be incorporated into the new SAKe designs: ten amino acids for type A SAKe (S6A) and six for type B (S6B)s.

Three different S6A (S6AE, S6AR and S6AC) and S6B (S6BE, S6BR and S6BC) scaffolds were selected for experimental evaluation. The appended letters refer to the selection criterium used: Rosetta Talaris2013 energy score (E),^43^ RMSD deviation from the idealized symmetric backbone architecture (R), or a combined rank score from criterium E and R (C). Analysis of the energy landscapes after RE_3_Volutionary design showed that the original Keap1 repeat sequences had worse scores compared to most inferred sequences.

All six proteins were successfully expressed and purified from *E. coli* BL21 (DE3) and circular dichroism (CD) spectroscopy showed spectra matching that of the parent template Keap1 β-propeller **(Figure S2)**. We identified a positive 233 nm CD signal, representing tertiary structure features, that was used to derive SAKe’s melting temperatures. The results show exceptional thermal stability, with S6BE exhibiting a melting temperature (T_m_) of over 95 °C. This is significantly higher than the melting temperature of the template Keap1 protein (T_m_ of 44.1 ° C) **(Figure 1)**.

The methodology used for the design of the SAKe backbone relies on ancestral sequence reconstruction. This method is known for yielding very stable proteins as it biases the sequence reconstruction on structurally relevant amino acids. Repetition of such sequences into a symmetric globular protein explains the increase in thermal stability of the different SAKe scaffolds respectively to their natural counterpart (Keap1).^28,44–47^

In order to confirm correct folding of the SAKe proteins, all purified proteins were subjected to crystallography. All proteins crystallized within days, diffracted with a resolution ranging from 1.3 to 1.95 Å and were successfully phased with their designer templates, confirming in this way the accuracy of the initial predictions **(Figure S3)**.

### SAKe loop variability

Loop modifiability is necessary for designing a protein binding interface, for example antibody-like protein binders, and for later functionalization of protein-based surface assemblies.^48^ To assess loop modifiability, three loop variants of the two most stable C_6_ symmetry SAKes: S6AC and S6BE were created.

The L1 loop was created by conserving the intersection of S6A-type and S6B-type loops, while adding four amino acids. For L2, a larger part of the S6A-type loop was conserved with the insertion of four amino acids. In both cases, these inserted four AA were randomly generated following the amino acid occurrence derived from a database of known nanobody CDR motifs.^49^ Only sequences containing at least one tyrosine and histidine were accepted, as these residues are often found to mediate interactions between antibody and antigen,^50^ while symmetric histidine arrangements were used before to scaffold metals and metaloxo clusters.^29,31,51^ The resultant sequence motifs are not commonly found in the natural human Kelch proteins. L3 describes a 3-fold symmetric variant with two long loops of various lengths: The longest loop in the L3 configuration was extracted from the crystal structure of the Kelch domain of human KBTBD5.^52^ S6AC/S6BE-L1, S6AC/S6BE-L2 and S6AC/S6BE-L3 have loop lengths of 9, 12 and alternating 10-15 amino acids, respectively. Finally, to assess the influence of loop composition on protein stability, we also transferred the original S6BE loop to S6AC, yielding S6AC-LB **(Table S1**). All new variants were successfully purified, and the structures of S6BE loop variants and S6AC-LB were confirmed via X-ray diffraction following an identical approach described above **(Figure 1)**.

With a T_m_ of 51.7 °C, the SAKe with the longest loops (10-15 AA) is still more stable than the natural Keap1 β-propeller (4-9 AA loops, T_m_ of 44.1 °C) **(Figure 1)**. The shortest loop (6AA) shows a Tm exceeding 95 °C and longer loops progressively lower thermostability. Interestingly, S6AC seemed the most moldable scaffold, retaining high stability even when carrying the loops which significantly destabilized S6BE **(Figure S4)**. Hence, S6AC core proved to accommodate a larger range of loops lengths. S6AC was designed from blade 1 of the Keap1 protein with a loop length of ten amino acids unlike S6BE which initial blades contained only six amino acids. This small difference biased the S6AC core to accept longer loop sequences.

For the continuation of this work we however continued with S6BE as S6AC derivatives precipitated in a buffer screening experiment at even mildly acidic pH, whereas the S6BE appeared to crystallize spontaneously.

### Engineering of the self-assembling SAKe

While protein crystallization cannot directly point towards protein self-assembly, there is a strong interplay between self-assembling forces and the crystallization process.^53^ Particles that self-assemble deliver faster crystal growth as the energy barrier for nucleation is substantially reduced.^54^ This idea was exploited to engineer new protein-protein and protein-surface interfaces for S6BE to improve self-assembly on mica surfaces.

As a first step to engineer a SAKe protein capable of self-assembling on the mica surface, the PPIs needed to be understood to find key residues stabilizing such a process. For this, S6BE was submitted to a pH titration experiment which highlighted the importance of neutralizing the protein charge to trigger spontaneous self-assembly. The high order of symmetry of S6BE together with its high pH stability led to spontaneous assembly of mm sized crystals in mildly acidic conditions (pH < pI), disassembling only at 1 pH point above the pI of the protein **(Figure 2)**. The self-assembled crystals showed a hydrogen bonding network stabilizing lateral contacts (growth on the longitudinal direction) and vertical contacts (stacking proteins one on top of another). The charge residues at the bottom loops (asparagine 30 and aspartic acid 31) stabilized the PPIs laterally interacting with the backbone of the neighboring protein. At the same time, proteins stack on top of each other stabilized by the tyrosine 57 at the top loops and the arginine at the bottom loops of the first blade (and equivalent for the other five blades) **(Figure 2)**. This led to the hypothesis that introduction of residues favoring surface-based assembly while preventing this vertical stacking of proteins would favor the formation of monolayers, by reducing the chances of having a multi-layered assembly or an on-surface adsorbed crystal.

**Figure 2:**
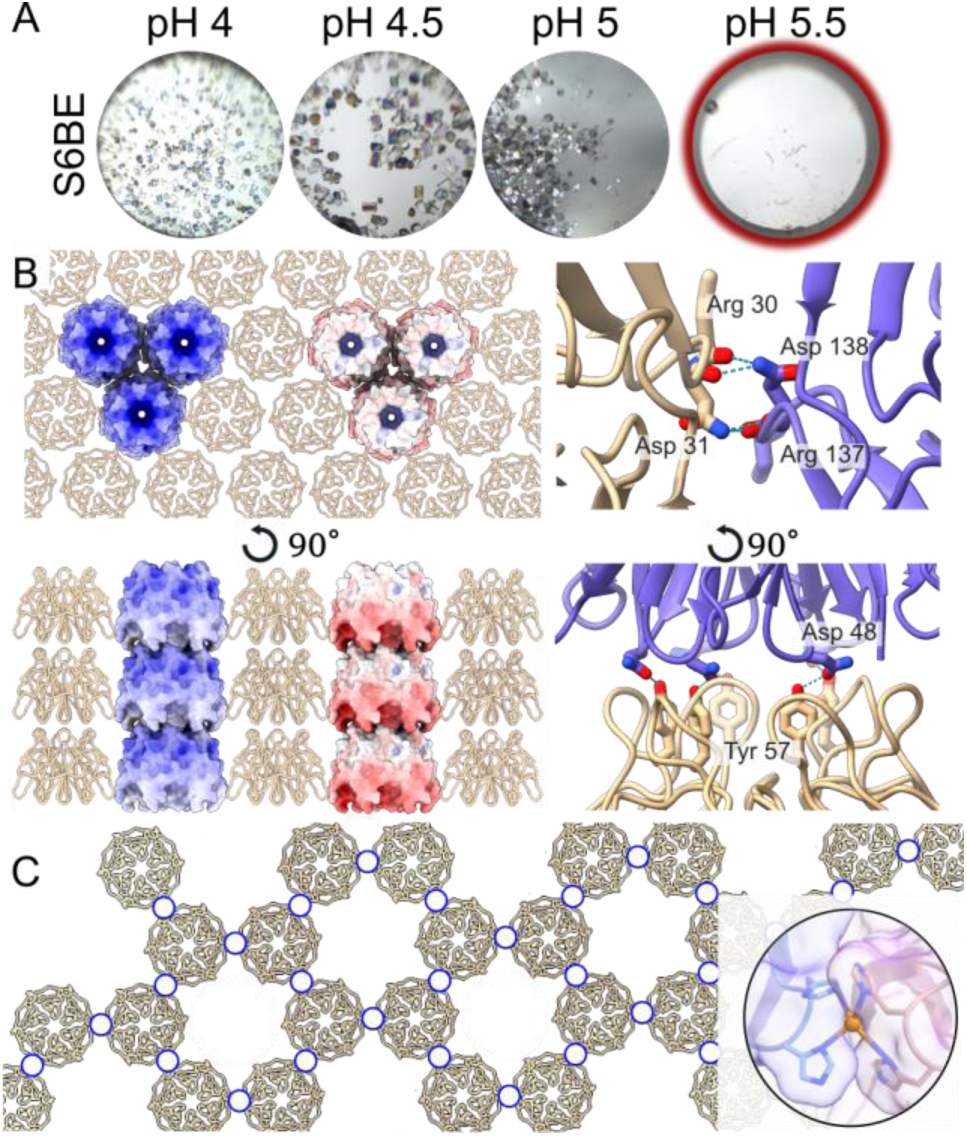
Design of the self-assembling variants: Dialysis experiments showcasing the pH sensitivity of S6BE crystallization and the pH sensitivity of the assemblies **(A)**. Similarly to (7ONC) a P1 symmetry was obtained, where proteins form hydrogen bonding laterally but also vertically. The electrostatic surface at pH 4 (left) and pH 5.5 (right) are displayed **(B)**. Based on the contact points of the S6BE crystals, 4 different SA variants were designed. Histidine residues were introduced in the 2^nd^, 4^th^ and 6^th^ blade or in all blades **(C)**. Schematic representation of the p3 intended monolayer 2D lattice. When the crystals formed at pH 4 were studied, a C121 symmetry was obtained, where proteins interacted laterally similar as in 7ONE, but also vertically, this time through the top loops.

Histidines are very versatile amino acids and are often used to drive self-assembly by metal coordination through the protonated π nitrogen.^55–58^ Another great trait of histidine residues is the imidazole ring, which behaves as an aromatic ring and is known to interact via π - π stacking with surrounding imidazole rings.^59–61^ These two interaction strategies are of great advantage when designing a self-assembling system. In the case of the π - π stacking not being strong enough to drive the self-assembly, metals could be introduced in the system to trigger the coordination and hence ease the formation of nucleation points. Therefore, two histidine residues were placed in exchange of the glutamic acid and the arginine residue creating a bis-his clamp to coordinate Zinc cations in order to gain control over the self-assembling process of the SAKe protein **(Figure 2)**.

Histidine residues were introduced by stepwise substitution of the residues involved in the hydrogen bonding with the backbone of the neighboring proteins stabilizing the lateral interaction (R30 and E31 and subsequent residues for each blade). S6BE-3HH (7OPA) with substitutions at the bottom protrusions of the 2nd (E76H and R77H), 4th (E170H and R171H) and 6th (E264H and R265H) blades (C3 rotational symmetry), and S6BE-6HH where the substitutions were done at the bottom protrusions of all the blades (C6 rotational symmetry) were designed to optimize the interaction between the proteins and the mica surface **(Figure S6)**. As a control to investigate whether both histidine residues are required for the self-assembly on the mica surface of the SAKe protein, two intermediate mutants were created, where the blades 2^nd^, 4^th^ and 6^th^ were mutated back either the first histidine residues (S6BE-3EH) or the second histidine residues (S6BE-3HR).

The contact points of the S6BE crystals were of electrostatic nature as the protein-protein interface was stabilized by hydrogen bonds, indicating a possible charge dependency **(Figure 2)**. In order to test such hypothesis, the in-solution self-assembling properties of the new variants (S6BE-3HH, S6BE-6HH, S6BE-3EH and S6BE-3HR) and S6BE were tested through dialysis experiments conducted at various pH values. The pH at which S6BE and S6BE-3HH assembled was found to be around 4 and 4.5 for S6BE-6HH. Notably, S6BE-3HH crystals required a week to grow, while the others assembled within a day **(Figure S5)**. This delay in crystal growth could be attributed to the loss of the mrotational symmetry and hence a higher entropy change upon assembly.^62^ The S6BE crystals remained stable up to pH 5 while the crystals of S6BE-6HH remained up to pH 7. This difference on the disassembling pH could be understood from two fronts. First, the isoelectric point of S6BE-6HH is one point higher than that of S6BE. In a system that is highly controlled by the electrostatics of the monomers, maintaining charge neutrality increases the likelihood of monomers interacting in a stable manner. Second, replacing two charged residues (the asparagine and glutamic acid) at the sides of the proteins by histidines which remain neutral at pH 7, reduces the electrostatic repulsion and increases the pH of disassembly.^63^

Still, both proteins readily reassembled upon lowering the pH again, highlighting the reversibility. To study the interactions driving the self-assembly, the crystals were analyzed using X-ray diffraction **(Figure S11)**. The packing arrangement was nearly identical to the one obtained with vapor diffusion at pH 7.0 and at pH 8.0 for S6BE (PDB 7ONE).

The symmetry and dipole-like character of S6BE along with the shape complementarity likely facilitated self-assembly into a tightly packed hexagonal structure where a network of hydrogen bonds and hydrophobic interactions further stabilizes the assembly. As pH increases above the pI of the proteins, residues Asp11, Asp23, Glu20, Glu28 and His15 (and the equivalent for the other blades) will lose the protonation state and the negative net charge of the protein is restored, triggering disassembly.

One of the engineered contact points was a hydrogen bond between the arginine residue and the backbone of the neighboring protein. This hydrogen bond was expected to not be altered by the change of pH as the pKa of the side chain of the arginine residue is around 10. Hence, in order to test whether both residues, the glutamic acid and the arginine were required to be mutated off, two intermediate SAKe mutants were designed, S6BE-3EH and S6BE-3HR **(Figure S6)**. These new mutants were submitted to the dialysis experiment as well and followed a similar trend as their parent protein. This time, the crystals could only be visualized after 48h incubation at pH 4. These crystals, for either protein, dissolved at pH 5.5 agreeing with the hypothesis of a pH driven crystallization **(Figure S5**). These results indicate that the high order symmetry of the SAKe scaffold is sufficient for the formation of the crystals. The addition of double histidine moieties in each blade substantially increases the stability of these crystals and renders them less sensitive to pH variations. Moreover, their introduction was expected to impact the interaction between the protein and the mica surface, having a key effect on stabilizing the assemblies on the surface.

### Metal-induced self-assembly

Metal-mediated protein self-assembly is not a new technique. It is commonly found in nature^55,56^ and has been used by many to engineer new PPIs points in protein building blocks. Working with a symmetric protein it is theorized to bring an advantage when it comes to engineering these new metal-mediating points, as it substantially reduces the search space for designing metal coordination points.

Metals offer a list of benefits when compared to noncovalent interactions. Metal coordination bonds are deemed to be of stronger nature and highly controllable when working with metals like Zn^2+^. Moreover, they offer a certain degree of directionality implied by the coordination geometry preferred by the metal, which is advantageous when designing protein assemblies.

Bis-his clamps were introduced in order to coordinate Zn^2+^ with the purpose of forming open honeycombs on the mica surface (S6BE-3HH) **(Figure 2)**. The effect of adding zinc cations was first studied by dynamic light scattering (DLS) to assess how long and at which ratios bigger molecules would form **(Figure S7)**. As metal coordination is pH-sensitive, three different values were used in these experiments, pH 5, 6 and 7 **(Figure S7)**. These values were chosen as beneath pH 5 already led to the formation of crystals in solution, so the appearance of peaks representing larger complexes would not indicate a direct effect of metal coordination. Another reason is that the histidine residues need to be neutral in order to coordinate metal cations.^64,65^ When zinc nitrate was added in the solution, there was a clear formation of large complexes already after 10 minutes incubation time. It is worth mentioning that at 1:1 ratio, a large fraction of protein remained in the monomer state, most likely due to insufficient number of metal cations to interact with the proteins. S6BE-3HH has, potentially, three coordination points per protein. At a ratio of 1:1, not all these coordination points will be occupied. At excess of metal cations, it is more clear how proteins start to easily coordinate these metals and form what could be protein aggregates or protein assemblies. To investigate whether these macrostructures were ordered, the 1:20 excess metal condition was visualized in the AFM **(Figure S8)**. A protein concentration of 16 µM led to direct aggregation and so, the protein concentration was reduced to 1 µM to perform the analysis. After 20 minutes of incubation (the time required to prepare the solution and to calibrate the AFM parameters), the surface exhibited no discernible order, indicating the formation of protein aggregates rather than protein assemblies. Interestingly, the control experiment led to the formation of ordered assemblies on the surface, what prompted the hypothesis that metal coordination was not necessary for the self-assembling of the SAKe proteins.

### On surface self-assembly

On-surface assembly on mica was investigated through amplitude-modulated atomic force microscopy (AFM). Since pH was anticipated to act as the self-assembly trigger, protein deposition was carried out using solutions spanning a pH range from 4 to 7.

S6BE showed a large number of proteins adsorbed onto the surface at pH 4, although no ordered assemblies were observed. At pH 7, no adsorbed protein was detected, most likely due to electrostatic repulsion between the negatively charged protein and the negatively charged mica substrate.

In contrast, S6BE-3HH and S6BE-6HH exhibited enhanced on-surface self-assembly. Unlike S6BE, both variants formed assemblies on the substrate, though with distinct features.

For S6BE-3HH, self-assembled arrays were observed across all tested pH values except pH 6, which is slightly above the isoelectric points of S6BE-3HH (5.32). Notably, the arrays observed at pH 7 exhibited more uniformity, with most proteins oriented along their vertical axes, an interaction that is showcased by the fiber-like structures imaged at the surface. At pH 4 and 5, the arrays imaged did not exhibit long-range order nor uniformity. This diversity on morphologies highlights the critical role of electrostatic interactions. In the case of pH 7 favoring radial over tangential growth, while at pH 4 and 5 there is no clear preference of growth (**Figure 4)**. As discussed for S6BE, the protonation state of the protein governs the strength of the interaction between the proteins and the mica surface. Protonation of N^π^ and N^τ^ nitrogens is expected to stabilize the protein on the mica surface by interacting with the negatively charged hydroxyl groups. With increasing pH, deprotonation of these nitrogens weakens the protein–surface interaction, thereby limiting the formation or stabilization of ordered assemblies on the surface. The necessity of this double histidine moiety for the formation of assemblies is showcased by the lack of assemblies found on the mica surface for S6BE-3EH and S6BE-3HR **(Figure S9).** These two proteins failed to form assemblies on the surface while crystals successfully formed on solution at pH 4. S6BE-3EH does not have the first histidine mutation which is expected to point towards the mica interface, stabilizing the proteins and the assemblies on the surface. S6BE-3HR on the other hand has a relatively large and flexible residue at the interface between neighboring proteins, destabilizing the PPIs and hence reducing the stability of the assemblies on the surface as the degrees-of-freedom of the proteins are reduced.

**Figure 4:**
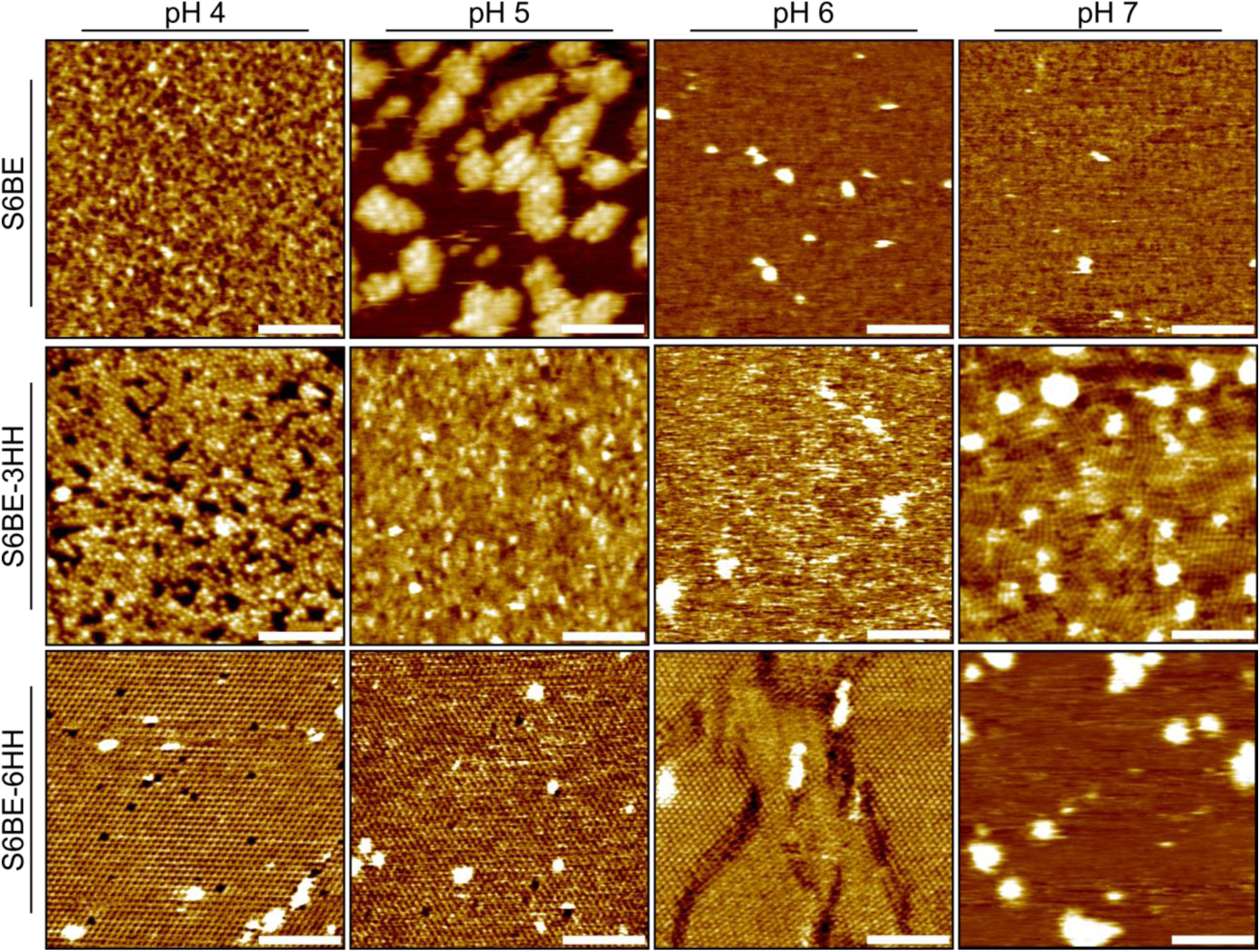
SAKe self-assembly on muscovite mica: Overview of AFM topography images acquired on muscovite mica for S6BE, S6BE-3HH, and S6BE-6HH at progressively increasing pH values (pH 4 – pH 7), highlighting the critical role of the presence and protonation state of the incorporated histidine residues in governing on-surface assembly. The scale bar is 50 nm length.

In contrast to S6BE-3HH, S6BE-6HH formed extended hexagonal assemblies at pH 4 and 5 that were readily visualized and resulted in high surface coverage. Upon increasing the pH to 6, these assemblies began to deteriorate and completely disappeared at pH 7 **(Figure 4)**. The observed structures followed the packing geometry anticipated from the design of the self-assembling variants **(Figure 5).** The proteins most likely anchor to the mica surface via their bottom loops, while lateral interactions between the protein sides establish the contact points that stabilize the lattice. This arrangement would promote on-surface assembly with the top loops exposed to the solvent, rendering them accessible for subsequent functionalization **(Figure 5)**. To achieve sub-monolayer coverage of the mica surface, the protein concentration was reduced to 0.01 uM, which is 100-fold lower than the working concentrations used for S6BE and S6BE-3HH, further highlighting the robustness and stability of the S6BE-6HH assemblies on the surface **(Figure S10)**.

**Figure 5:**
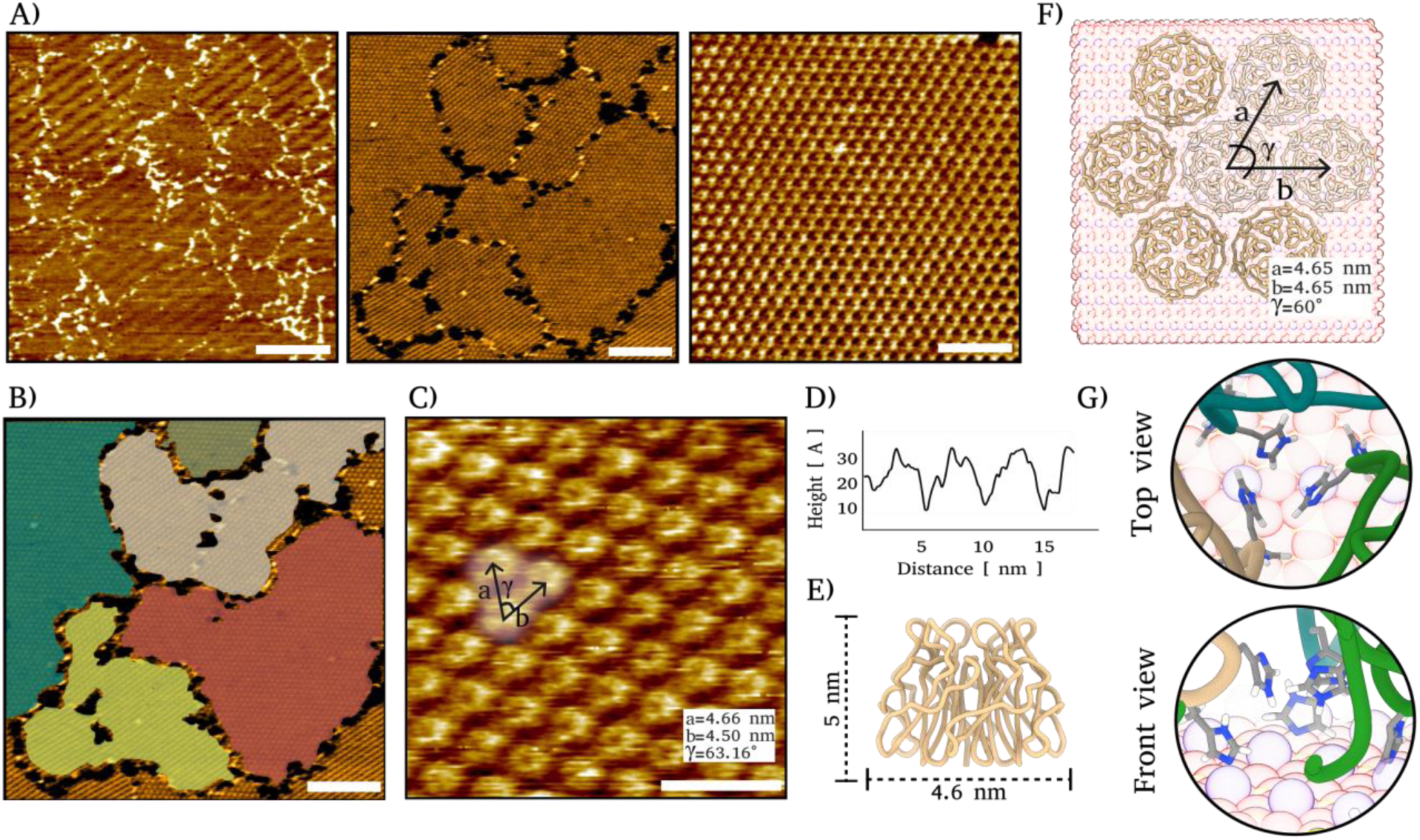
High resolution imaging and assembly mechanism on Mica: High resolution images of assemblies of S6BE-6HH on muscovite mica showcasing the vast surface coverage as well as the lattice periodicity. The scale bars in the figure represent from left to right 200, 50 and 20 nm respectively **(A)**. Independent domains were consistently found on the surface arising from different directions of growth. Scale bar represents 50 nm **(B)**. Analysis of the lattice periodicity shows unit vectors characteristic of C6 symmetry, with a repeat distance corresponding to the dimension of a single protein. These experimentally derived vectors are in agreement with those predicted by the structural model **(C-F)**. A schematic representation illustrates the proposed interaction between the mica surface and the histidine residues located in the bottom loops of S6BE-6HH, highlighting their role in anchoring and stabilizing the on-surface assembly **(G)**.

Interestingly, the on-surface assemblies do not fully mirror the trends observed in solution. Although increasing the pH beyond the protein’s pI markedly affects the overall charge, producing pronounced effects for S6BE and S6BE-6HH, the loss of protein–surface interactions, and thus on-surface self-assembly, occurs at lower pH values than those required to disrupt assembly in solution. With respect to assembly kinetics, no discernible differences were observed between S6BE-3HH and S6BE-6HH. For both variants, assemblies formed within the first 10 minutes (corresponding to the time needed for AFM cantilever calibration) and remained stable throughout imaging. However, the two-dimensional assemblies formed by S6BE-3HH do not replicate the crystalline lattice observed in its self-assembled crystals, whereas those formed by S6BE-6HH closely resemble the lattice architecture obtained in the crystalline state **(Figure S11, S12)**.

As only single-layer assemblies are observed, with no evidence of multilayered or overlaid crystal precipitation, it’s unlikely that these on-surface assemblies are the result of the precipitation of preformed crystals in solution **(Figure S12)**. Moreover, if crystal precipitation was the dominant mechanism, S6BE would be expected to form large assemblies, which is inconsistent with the experimental observations **(Figure 4)**.

The proposed mechanism underlying SAKe protein self-assembly extends beyond protein–protein interactions (PPIs) and incorporates a crucial contribution from protein–surface interactions. Muscovite mica becomes negatively charged upon immersion in aqueous solution due to the desorption of interfacial potassium ions. During overnight incubation, small protein nuclei are expected to form in solution; these nuclei subsequently precipitate and interact with the mica surface. This interaction is likely mediated by the histidine residues engineered on the bottom face of the protein **(Figure S6)**. Only under conditions where the histidines remain protonated favor (i) stable anchoring these nuclei to the surface and (ii) continued adsorption of monomers from the bulk solution, followed by lateral assembly growth **(Figure 4)**. Because nucleation occurs randomly across the surface, growing arrays expand outward until they encounter neighboring domains. When two misaligned arrays meet, a domain boundary is formed **(Figure S13)**. The templating influence of the surface is further evidenced by the angular relationship between adjacent domains. A consistent angle of 160° ± 5° is observed, reflecting the pseudo-C6 symmetry imposed by the mica lattice **(Figure 5)**. Additional support for a surface-mediated assembly mechanism is provided by the clear differences in packing between the S6BE-3HH crystal structure **(Figure S11)** and the smaller surface-confined arrays **(Figure S12)**, indicating that the mica substrate modulates the packing orientation of the protein.

## CONCLUSION

Our design strategy highlights the value of using pseudo-symmetric natural scaffolds for the design of lattice forming building blocks. The variability of the Kelch fold implies a wide variety of loop motifs are supported while the bottom remains modifiable for surface interaction. High-resolution AFM confirmed that addition of surface-facing histidines successfully formed ordered arrays on muscovite mica (up to 5 µm^2^). The intrinsic symmetry of the SAKe protein together with its charge distribution strongly contributed to the stabilization and the in-solution and on-surface morphology of the assemblies. The additional modifiability of this scaffold makes it a valuable building block for technologies such as biosensing or on-surface catalysis.

## Acknowledgements

We thank Prof. Tatjana P. Vogt for granting us use of the CD spectrometer. We thank the beamline scientists at Diamond Light Source, Swiss Light Source, European Synchrotron Radiation Facility and Elettra Synchrotron and their scientists for their assistance. This work was supported by Research Foundation Flanders (FWO) (1S89918N, G0F9316N and G051917N, ZKE-1919-04-W01) and by KU Leuven-Internal Funds (C14/23/090).

## Materials & Methods

### Computational design

SAKe was designed using the RE_3_Volutionary protein design method^28^ using the KELCH domain of human Keap1 (PDB code: 1ZGK) as a template.^33^ Rosetta symmetry docking was used to construct C_6_ symmetric backbone models, using blade structures extracted from the human Keap1 β-propeller.^41,66^ We used the 2^nd^ blade for backbone modeling of A-type SAKe, and the 6^th^ blade to design the B type SAKe backbone. For each run, 20000 backbone models were generated. The models were filtered on Rosetta symmetry docking energy scores and RMSD from a manually constructed symmetric backbone. Clustal Omega was used to generate multiple sequence alignments (MSAs) from the six repeats of the Keap1 β-propeller^67^. When designing B type SAKe, the conserved VAPM motive was erroneously replaced by VAPL during MSA. The MSAs and their accompanying unrooted phylogenetic trees were used to construct lists of putative ancestral sequences using the FastML server,^68^ with 250 sequences per node for a total of 1000 sequences for each SAKe construct. The ancestral sequences were repeated 6-fold to fit the full length of the symmetric backbones. The putative ancestral sequences were mapped on their corresponding backbone models using a custom PyRosetta script, which was written by prof. D. Simoncini and is available on our laboratory GitHub repository (https://github.com/kullbmd/kullbmd).^28^ Talaris2013 energy scores and RMSD calculated against the input backbone model were used to filter for viable designs.^43^ For all SAKe constructs except S6AR, cysteine residues were then mutated to serine or alanine, depending on their locations.

### Randomized loop generation

First, a Python2.7 script was used to calculate the percentage of occurrence for each amino acid in the iCAN database file, which contains various nanobody CDR motifs.^49^ As input, we provided a TXT file containing all iCAN database sequence strings. Then, another Python2.7 script was used to randomly generate ‘b’ number of sequences with length ‘l’. The script uses the previously determined percentages of occurrence and selects only sequences which contain both a Tyr and a His. These scripts are available in the supplementary information **(Supplementary Scripts)**.

### Cloning

Amino acid sequences were first reverse translated into DNA sequences, using a codon optimization tool by IDT (Integrated DNA Technologies, Haasrode, Belgium). DNA was ordered as gBlocks from IDT and subsequently subcloned in pET-28a(+) via NdeI and XhoI restriction sites. Alternatively, DNA was pre-cloned in pET-28a(+) plasmids by BGI (BGI genomics, China). Primers were bought from IDT. All PCR reactions followed the Phusion^TM^ High-Fidelity DNA Polymerase protocol (Thermo Fisher Scientific, Massachusetts, United States). All restriction enzymes were bought from ThermoFisher (FastDigest product line). Ligations were performed with the T4 DNA Ligase from Promega (Promega, Wisconsin, United States). DNA was purified using the GeneJET Plasmid Miniprep Kit, GeneJET PCR Purification Kit and GeneJET Gel Extraction kit from Thermo Fisher Scientific. Before Sanger sequencing by LGC (LGC genomics, Teddington, United Kingdom), recombinant vectors were transformed to chemically competent *E. coli* DH5α using standard heat shock transformation. For protein expression, DNA isolated from single colonies was transformed to chemically competent *E. coli* BL21 (DE3) via heat shock transformation.

### Protein expression

*E. coli* BL21 (DE3) transformed with protein-encoding pET28 plasmids, were grown in 1 L LB (with 0.05 mg/mL kanamycin) at 37 °C, while shaking. At an OD_600_ of 0.6, the cells were left to cool on ice for 20 min. After adding 0.5 mM IPTG incubation continued at 20 °C for 16-18 h while shaking. Cells were harvested via centrifugation at 6721 g. The supernatant was discarded, and pellets were immediately stored at −24 °C.

### Protein purification

**SAKe proteins:** Pellets were thawed on ice and suspended in lysis buffer (40 mL, 50 mM NaH_2_PO_4_, 200 mM NaCl, 10 mM imidazole, 1 mM phenylmethylsulfonyl fluoride (PMSF), 30 mg hen eggwhite lysozyme (HEWL), pH 8.0). They were then incubated for 30 min at 15 °C while rotating head-over-head. After, they were lysed via sonication. Lysates were centrifuged at 20216 g for 30 min. The Supernatant was filtered (0.45 µm) and loaded on a 5 mL Ni-NTA column equilibrated with buffer A (5 CV, 50 mM NaH_2_PO_4_, 200 mM NaCl, 10 mM imidazole, pH 8.0). The column was washed with buffer A (10 CV, 50 mM NaH_2_PO_4_, 200 mM NaCl, 10 mM imidazole, pH 8.0) and buffer B (10 CV, 50 mM NaH_2_PO_4_, 200 mM NaCl, 20 mM imidazole, pH 8.0) and the proteins eluted with buffer C (10 CV, 50 mM NaH_2_PO_4_, 200 mM NaCl, 300 mM imidazole, pH 8.0). SDS PAGE was used to verify presence of the proteins in the eluate fractions. The fractions containing the proteins of interest were collected and dialyzed overnight in phosphate buffer (50 mM NaH_2_PO_4_, 200 mM NaCl, pH 8.0), using 7000 kDa cut-off SnakeSkin^TM^ Dialysis tubing (Thermo Fischer Scientific). At the same time, histidine tags were removed via thrombin digestion (100 U). The dialyzed samples were subjected to an additional Ni-NTA chromatography step, where the proteins now appear in the flow through. Next, the proteins were concentrated via ultrafiltration and injected on a Superdex 200pg 16/600 or Superdex 75pg 16/600 column (Cytiva, Hoegaarden, Belgium) equilibrated with elution buffer. UV_280_ absorbance peaks were collected and concentrated via ultrafiltration to stocks of 20 mg/mL or more. These stock proteins were stored at 4 °C.

**S6BE, S6BE-3HH and S6BE-6HH for AFM:** were purified as described above with the exception that 5 mM EDTA was added before SEC. **Keap1:** Because of the solvent-exposed cysteine, 1 mM freshly prepared DTT is supplemented in every step. This includes the sample preparation steps for SDS PAGE and crystallization.

### Circular dichroism spectroscopy

CD spectroscopy was performed with a JASCO J-1500 spectrometer (JASCO Inc., Maryland, United States). To measure the CD spectra, protein samples were diluted (400 µL, 0.1 mg/mL) in phosphate buffer (20 mM mM NaH_2_PO_4_, pH 7.6). Ellipticity was measured at 20 °C from 260 nm to 200 nm, using 1 mm cuvettes. Five accumulations were averaged. To estimate melting temperatures, protein samples were first diluted (400 µL, 0.05 mg/mL) in phosphate buffer (20 mM mM NaH_2_PO_4_, pH 7.6). Then, ellipticity was measured from 260 nm to 200 nm, from 5 to 95 °C with intervals of 5 °C, using sealable 2 mm cuvettes. A Python script was used to plot the data as a heatmap. Three accumulations were averaged. For accurate determination of melting temperatures, samples were diluted (400 µL, 0.25 mg/mL) in phosphate buffer (20 mM mM NaH_2_PO_4_, pH 7.6). The signal at 233 nm was followed from 5 to 95 °C with intervals of 0.2 °C, using sealable 2 mm cuvettes. The data was analyzed with a Python script, which fits a Boltzmann-sigmoid equation and extracts the midpoint T_m_. The parameters are described in a previous publication by Mylemans *et al.*^38^

### Dynamic light scattering

Concentrated S6BE, S6BE-3HH, S6BE-3EH or S6BE-3HR (20 mg/mL, 20 mM HEPES, pH 8.0) were diluted with either MES (20 mM, pH 5.6) or MilliQ to a concentration of 1 mg/mL. Metal suspensions (Cu(NO_3_)_2_ or Zn(NO_3_)_2_, MilliQ) were titrated to achieve the desired ratio of protein:metal. Size measurements were obtained at 25 °C using a Zetasizer Nano ZS instrument (Malvern Panalytical, Malvern, United Kingdom) and quartz cuvette (ZEN2112). Data analysis was performed using the Zetasizer software 7.11 (Malvern Panalytical).

### Atomic force microscopy

Proteins for AFM experiments were obtained from stock solutions (in 20mM HEPES, 200 mM NaCl, pH8) with 50% glycerol and stored at −80 °C. The samples were dialyzed overnight in order to prepare them with the imaging buffer, and further diluted to the working concentration. The buffers for the experiments were prepared according to the working pH. Buffers at pH 4 and 5 were prepared with 20 mM Na-Acetate, equilibrating the pH with 100% acetate. Buffers at pH 6 were prepared with 20 mM MES (2-(N-morpholino)ethanesulfonic acid). The pH was equilibrated with 6N NaOH. And the buffer at pH 7 was prepared with 20 mM of HEPES (4-(2-hydroxyethyl)-1-piperazineethanesulfonic acid). The pH was equilibrated with 6N NaOH.

To prevent cross contamination, all the utensils required to perform the analysis were cleaned by five minutes of sonication in isopropanol, water MQ quality and 100% ethanol spectroscopy quality and dried with N_2_ or Argon gas. All the images were taken in a Cypher S AFM Microscope from Oxford Instruments Asylum Research with a liquid perfusion cell (Cypher ES) and Fast-scanning high-frequency silicon probes (FS-1500AUD) from Oxford Instruments. Excitation of the tip was done by blue Drive photothermal excitation.

AFM images were processed using Scanning Probe Image Processor (SPIP, Image Metrology ApS), and Gwyddion software.

### Protein crystallization

Crystals were grown via sitting-drop vapour diffusion using NeXtal Crystal Screening kits (Qiagen), SWISSCI MRC 96-well plates (Molecular Dimensions Inc) and a Crystal Gryphon robot (Art Robbins Instruments). For native crystallography droplets consisted of 0.3 µL buffer and 0.3 µL protein (10 mg/mL in 20 mM HEPES pH 8.0, 200 mM NaCl). Plates were incubated at 20 °C. Protein crystals were vitrified after single-step soaking. PEG 400 or glycerol were used as cryoprotectant. X-ray diffraction experiments were performed at Diamond Light Source (United Kingdom), European Synchrotron Radiation Facility (France), Elletra Synchrotron (Italy) and Swiss Light Source (Switzerland).

### pH induced crystallization

To determine the pH tipping point of assembly of S6BE, S6BE-3HH and S6BE-6HH (5mg/mL), the three proteins were dialyzed at 20 °C in 500 mL of 50 mM citrate buffer at pH 4.0, pH 4.5, pH 5.0, pH 5.5, pH 6.0, pH 6.5; and in 500 mL of 20 mM HEPES buffer at pH 7.0, pH 7.5 an pH 8.0. To test the reversibility of the self-assembly of the three proteins, two experiments were carried out concurrently. One where the three proteins were dialyzed from pH 4.0 to pH 8.0, and one from pH 6.5 to pH 4.0. The latter would indicate the pH at which the crystals were readily visible. Such pH value (pH 4.0 for S6BE and S6BE-3HH and pH 4.5 for S6BE-6HH) was then used to dialyze the respective proteins upon dissolution of the crystals. For each protein, a self-assembled crystal was soaked in cryo-protectant, vitrified and shipped to Diamond Light Source (United Kingdom) and Elettra Synchrotron (Italy) for X-ray diffraction. Pictures of the self-assembled crystals were taken with a Nikon SMZ800N microscope, outfitted with a TV Lens C 0.45x (Nikon, Tokyo, Japan).

### Xray diffraction analysis

Diffraction patterns were indexed using XDS or DIALS.^69,70^ Data reduction was done with Aimless in CCP4.^71,72^ Resolution cut-offs were chosen to achieve an outer shell completeness of at least 92.5 %, while also satisfying the requirements of outer shell Rpim below 45 %, CC1/2 above 30 %, and I/sigma(I) above 1.0. Molecular Replacement phasing was done with PHASER, using our computationally designed models as search ensemble.^18^ Refinement was done manually with phenix.refine and Coot.^73,74^ The final structures were validated using Molprobity and the PDB validation tool,^75^ before being deposited to RCSB PDB.

### Determination of pI and surface electrostatics

Surface electrostatics were calculated at pH 4.0 and 8.0 via PDB2PQR, using a full-length SAKe as input. The calculated surfaces were visualized via PyMOL. pI values were calculated via PROPKA.^76–78^

