## Supplementary information for "Design and characterization of SAKe, a new building block for protein self-assembly"

**I. SUPPLEMENTARY FIGURES**

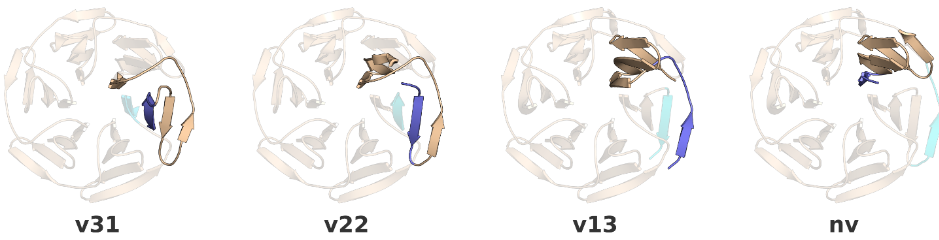

**Figure S1:** Velcro-closure in globular repeat proteins. A disparity between the structural and sequence-related repeat motifs can lead to so-called `velcro' closure of a protein fold. In the β-propeller example, a single repeat sequence forms this `velcro'-like closure when it spans two `blades': In v13 (velcro 1-3), the N terminal β-strand forms the last strand of one blade, while the three other β-strands of the repeat sequence are buried in the following/next blade. In nv (no velcro), the repeat sequence exactly matches a single `blade'.

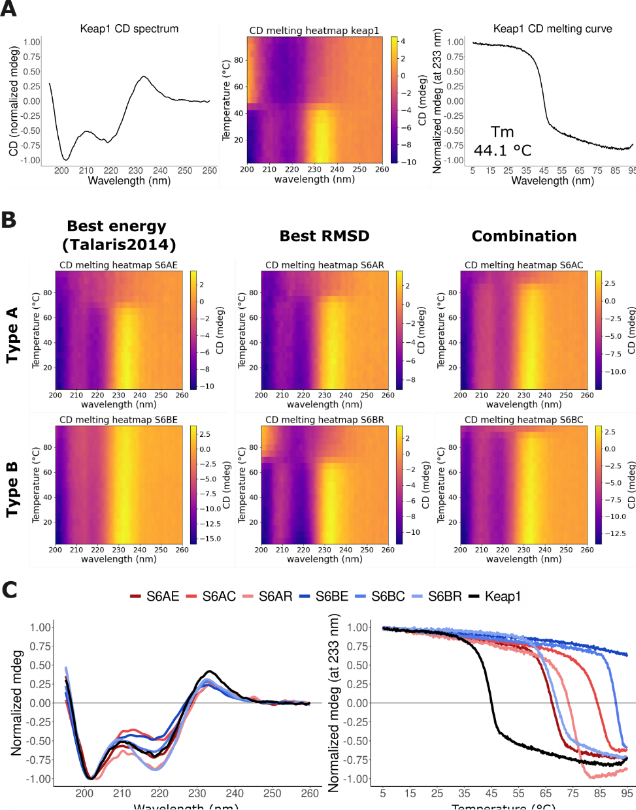

**Figure S2:** CD spectroscopy experiments on template protein Keap1 and the designer SAKe proteins. (A, left) CD spectrum of Keap1. The CD spectrum of Keap1 exhibited an unusual feature: A positive peak around 233 nm. This peak is a feature of the protein's 3D structure and can be attributed to the presence of aromatic amino acids such as tyrosine and tryptophan, which are known to contribute to near and far UV CD signals, for example as reported by Cantor and Timasheff in 1982 in their research on chymotrypsinogen activation, where they proposed that the CD signal at 230 nm is indicative of tertiary structure changes as the major alteration observed within this range could be attributed to conformational changes of one or more tryptophan residues.6 (A, middle) CD spectroscopy temperature interval analysis for Keap1. This heatmap plots the CD spectra measured from 5 to 95 °C at intervals of 5 °C. (A, right) Following the CD signal at 233 nm from 5 to 95 °C allows determination of the Keap1 melting temperature. (B) CD spectroscopy temperature interval analysis for S6C, S6D, S6E and S6F. These heatmaps plot the CD spectra measured from 5 to 95 °C at intervals of 5 °C. These heatmaps show that the 233 nm signal, which correlates to tertiary structure, can be used to more accurately follow SAKe melting. (C, left) CD spectra of all A and B type SAKe designs are like that of Keap1. (A, right) The corresponding melting curves were obtained via variable temperature measurement of the CD signal at 233 nm.

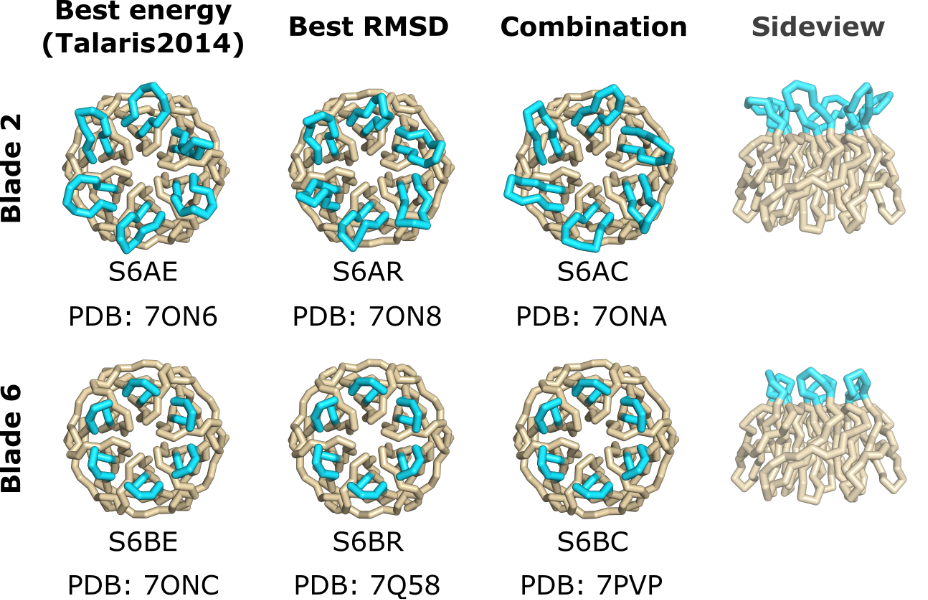

**Figure S3:** Summary of SAKe core designs. SAKe scaffolds were created by mapping ancestral sequences onto a symmetric backbone generated through Rosetta symmetry docking. This summary shows the specific blades from the Kelch domain of human Keap1 utilized in constructing the backbone and the scoring parameters (energy or RMSD) then employed to select candidates for experimental validation.

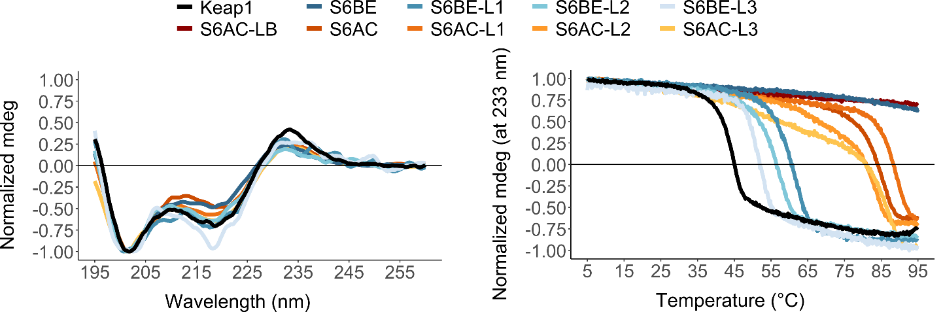
**Figure S4:** CD spectra and melting curves of SAKe loop variants. (Left) CD spectra of all SAKe loop variants are like that of Keap1. (Right) The corresponding melting curves were obtained via temperature interval measurement of the CD signal at 233 nm.

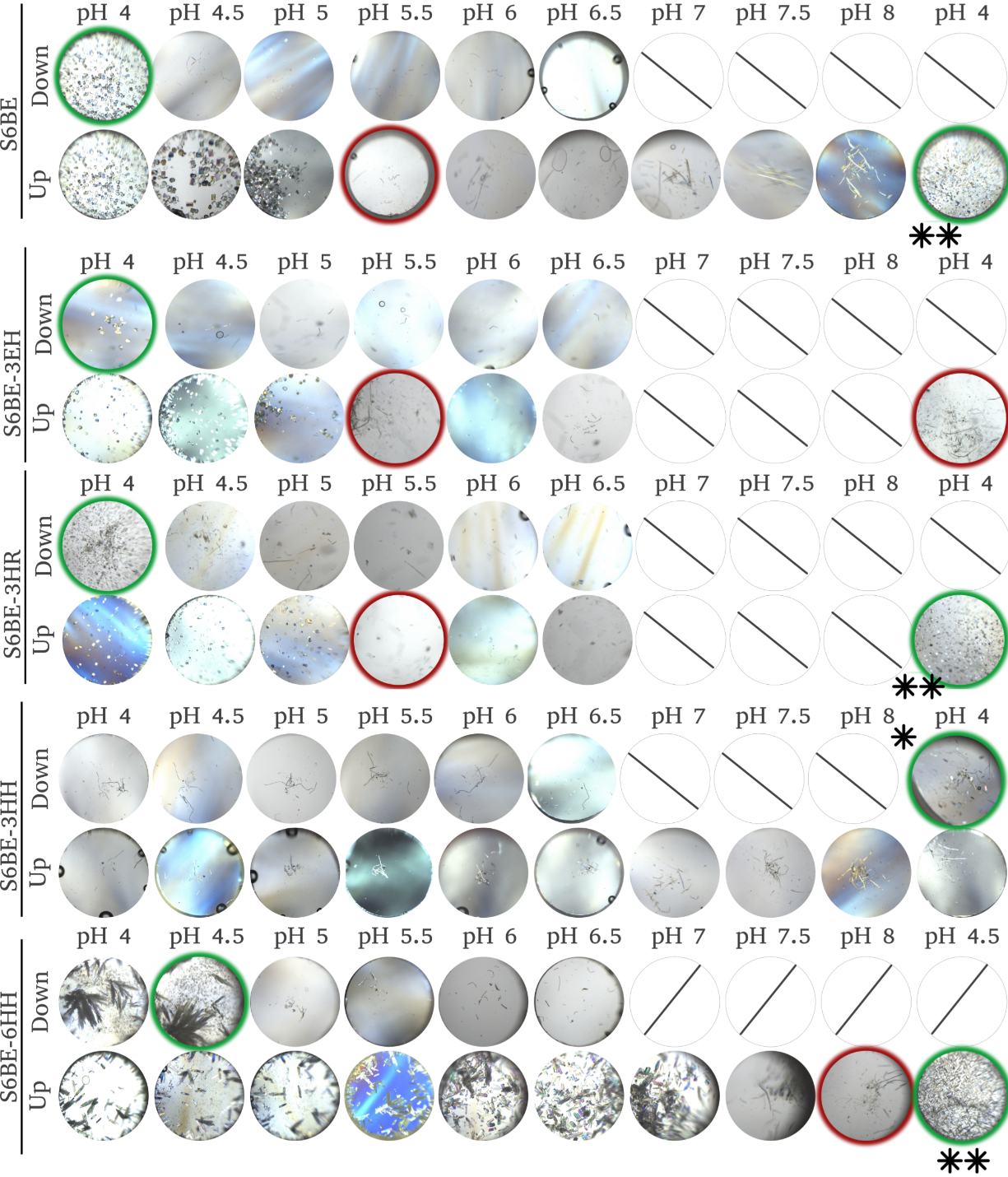

**Figure S5:** In-solution dialysis experiment: Images taken every 48h. All proteins were prepared at 5 mg/mL with 500 uL per dialysis cup. The proteins were dialyzed either from pH 4 to pH 8 (ascending pH) or from pH 6.5 to pH 4 (descending pH). Green circles indicate the respective pH at which crystals were first observed. Red circles highlight the pH at which these crystals were fully dissolved again. (*) S6BE-3HH crystals still form at pH 4.0 but require a full week for assembly. The reduced symmetry of S6BE-3HH likely leads to a much slower crystal assembly, due to a higher required change in entropy. (**) Following the complete dissolution of the crystals initially formed at pH 4, a 24-hour incubation at pH 4 (S6BE, S6BE-3EH and S6BE-3HR) or pH 4.5 (S6BE-6HH) again leads to in-solution crystal assembly, showing the reversibility of this process.

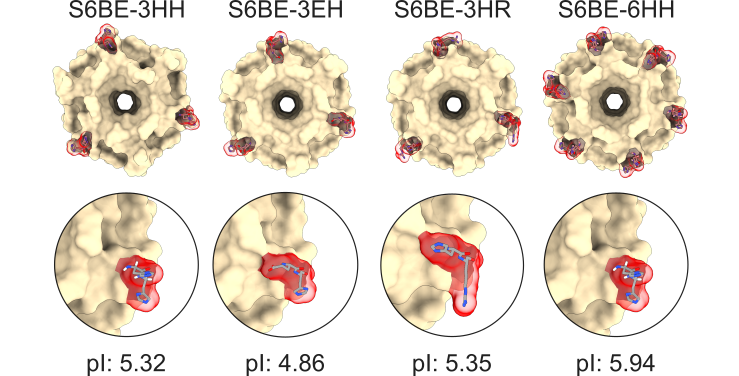

**Figure S6:** Self-assembling mutations of S6BE with their respective isoelectric points calculated by PropKa.

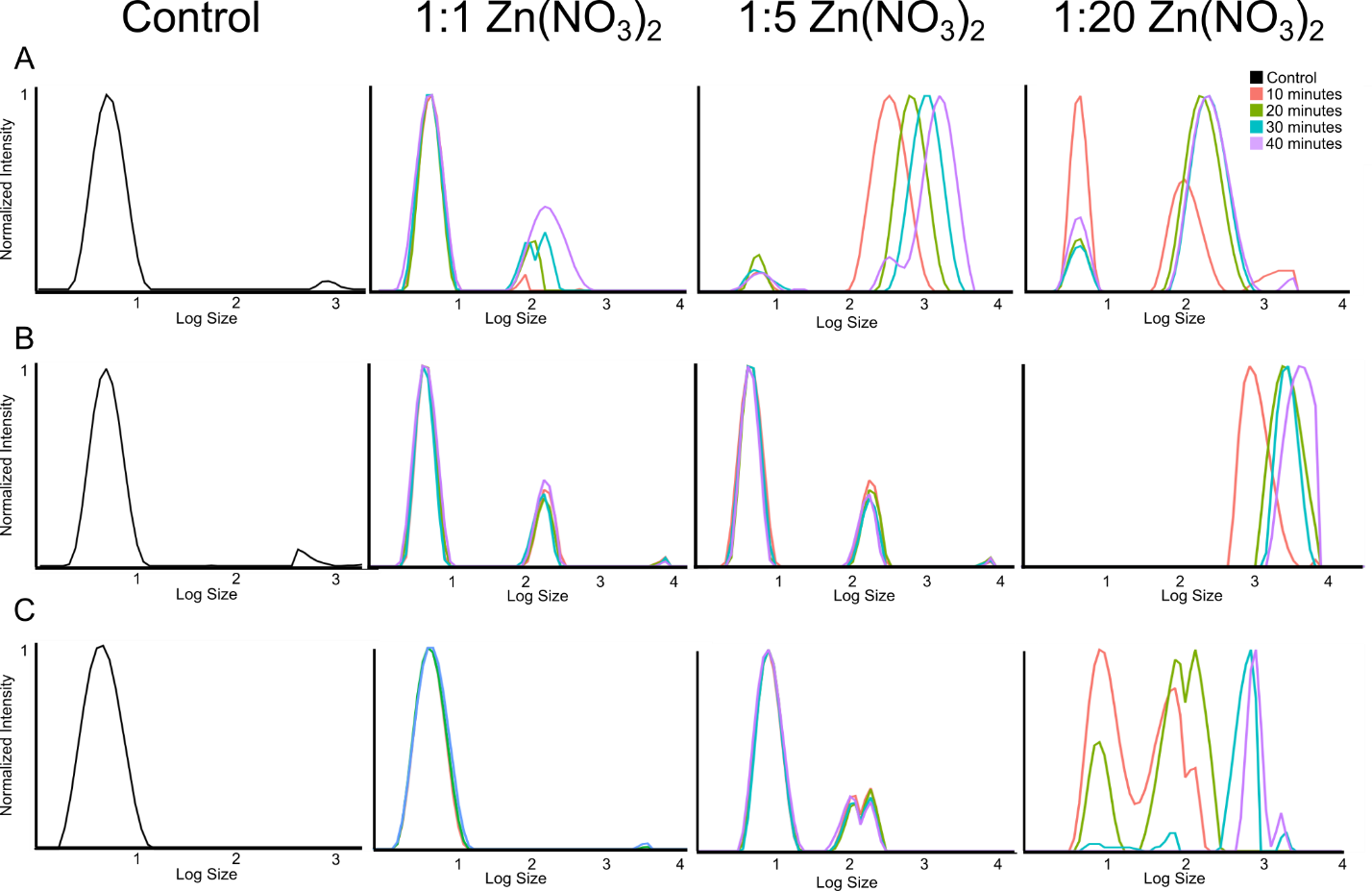

**Figure S7:** DLS spectra of S6BE-3HH with Zn(NO_3_)_2_: DLS spectra of the coordination of Zn^2+^ with the S6BE-3HH at different metal to protein ratio and at 20 mM MES buffer at pH 5, 6 and 7 (A, B and C respectively). Measurements were taken every 10 minutes to study the evolution of the system. In general, larger molecules are generated as the incubation time increases, seeing such effect clearer at ratios of 1:20 and at pH 7.

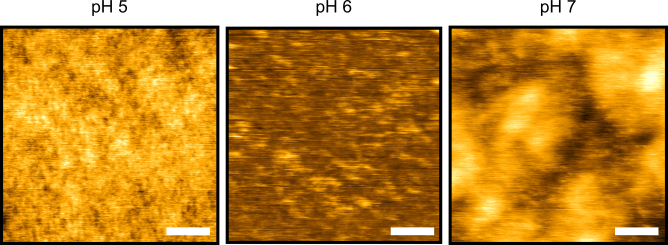

**Figure S8:** AFM images of S6BE-3HH with Zn(NO_3_)_2_: AFM imaging of the S6BE-3HH coordination of Zn(NO_3_)_2_ at a ratio of 1:20 on mica. Protein concentration was reduced to 1 uM due to the formation of aggregates at 16 uM. No assemblies could be observed. Instead, proteins were found to be without any order on the surface with some protein aggregates precipitated on the mica surface, agreeing with the observation in the DLS. The scale bar corresponds to 100 nm length.

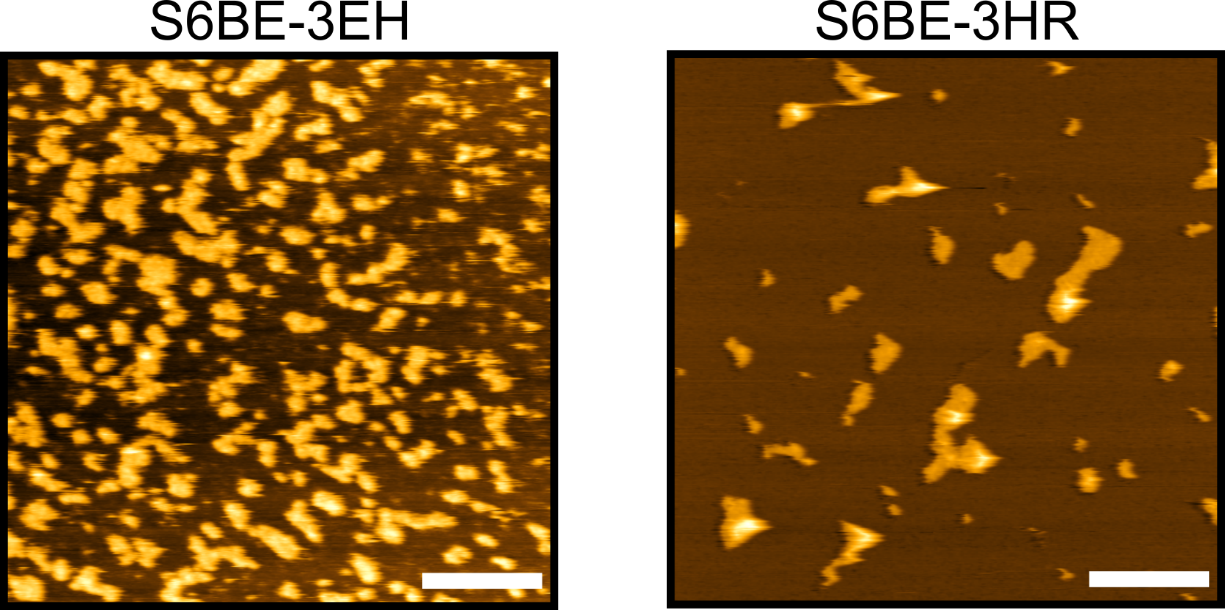

**Figure S9:** AFM images of S6BE-3EH and S6BE-3HR: Images were taken at 20 mM Sodium acetate buffer at pH 4 on muscovite mica. No assemblies could be found, and only small aggregates are imaged on the surface for either protein. The scale bar corresponds to 200 nm length.

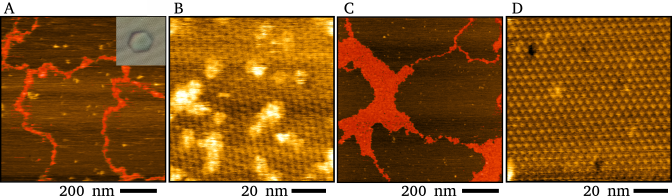

**Figure S10:** Concentration based AFM measurements of S6BE-6HH: Images taken of S6BE-6HH assemblies formed on the surface of mica at 1 uM (A,B) and at 0.1 uM (C,D). As can be observed the ten times dilution did not influence the formation of assemblies. Both images were taken with 20 mM Sodium Acetate buffer at pH 5.

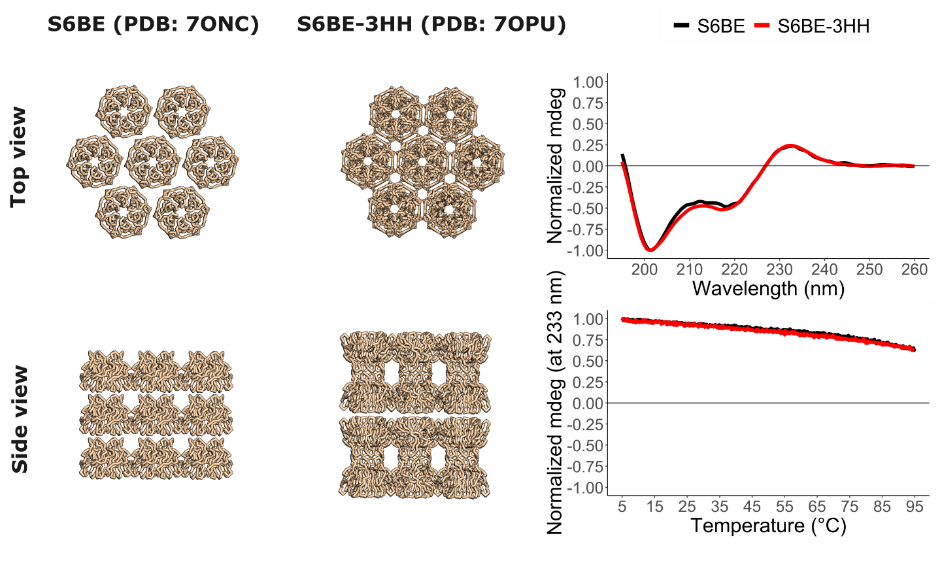

**Figure S11**: Crystal packing, CD spectrum and CD melting curve of S6BE and S6BE-3HH. Both proteins show similar CD spectra and melting curves. However, their crystal packing differs: S6BE-3HH prefers a packing arrangement where the proteins are organized in pairs of either top to top or bottom to bottom. In contrast, S6BE tends to favor a top to bottom arrangement.

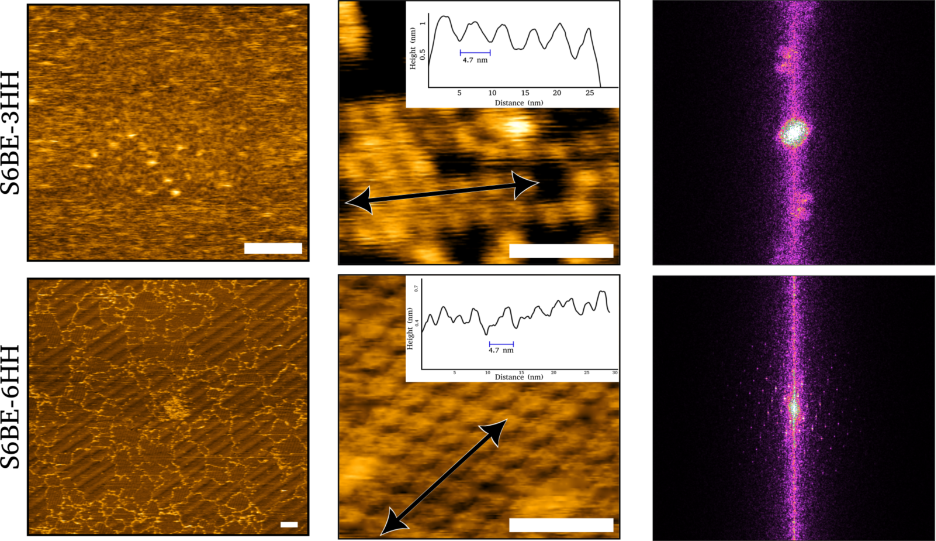

**Figure S12**: Profiles and 2D FFT analysis of the assemblies of S6BE-3HH and S6BE-6HH. The conditions used for imaging were 20 mM Sodium Acetate buffer at pH 4. Height profiles were taken across the observed assemblies. For S6BE-3HH assemblies could be found on a bottom later above which no ordered proteins could be found (top row). Despite observing some small ordered arrays, the 2D FFT analysis indicated no order along the full image. For S6BE-6HH (bottom row), an opposite scenario was observed. No indication of multilayer formation was found when large scale images were taken. High resolution images and their 2D FFT analysis indicated the formation of hexagonal assemblies on the surface.

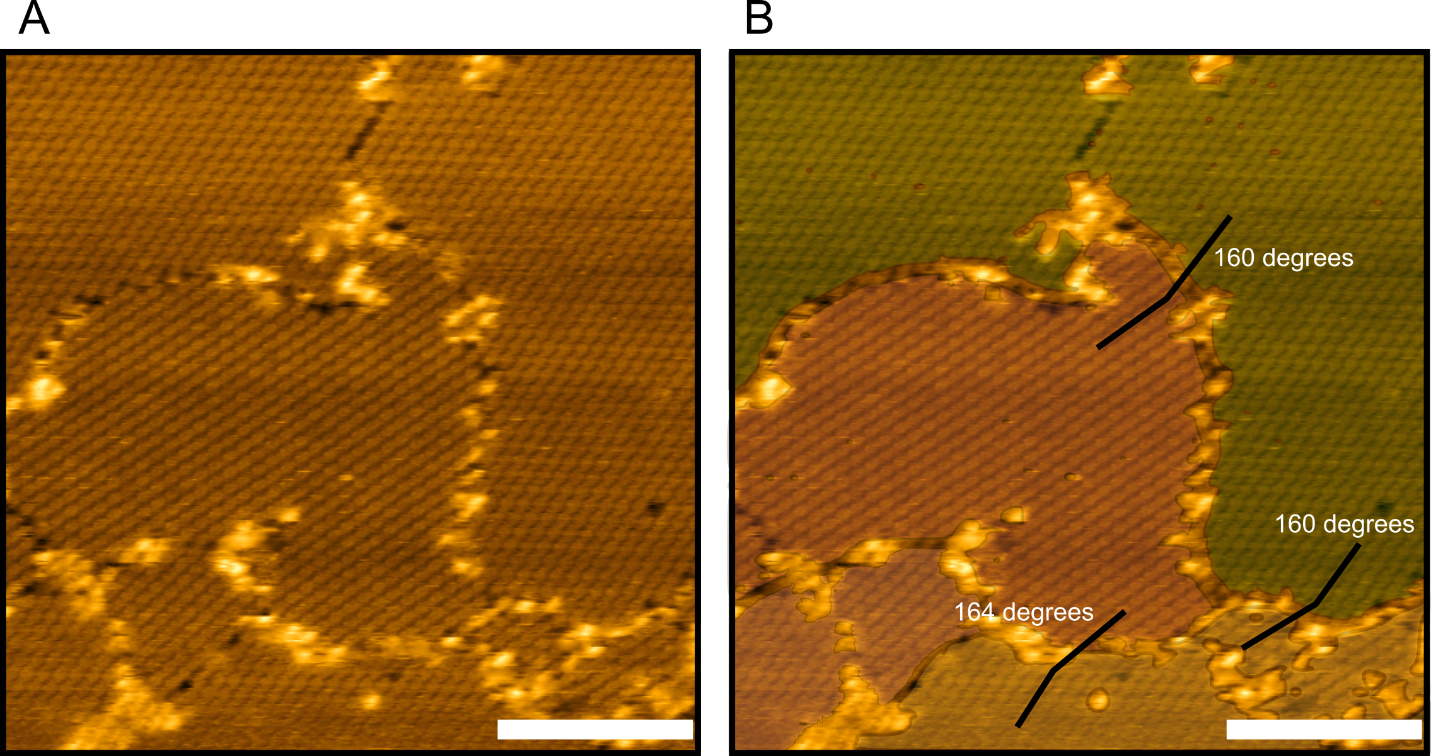

**Figure S13:** Domain difference analysis of S6BE-6HH assemblies: Images taken of S6BE-6HH with 20mM Sodium Acetate buffer at pH 4. The inlet bar corresponds to 50 µm. Domains have been highlighted with different colored masks B) and the angle between them was calculated with a profile line drawn from one domain to another in Fiji image processing software. 12 profiles were calculated yielding an angle of 161 +/- 5 degrees, following the expected deviation given the influence of the protein symmetry and the surface symmetry.

**II. SUPPLEMENTARY TABLES**

**Table S1:** Summary of SAKe melting temperatures, obtained via CD temperature interval analysis at 233 nm. Two melting temperatures indicated a two-state model was used to fit the data.

| Protein | PDB code(s) | MW (KDa) | Loop Length | Loop Sequence | Tm (°C) |
| --- | --- | --- | --- | --- | --- |
| Keap1 | 1ZGK | 34.02 | - | - | 44.1 |
| S6AE | 7ON6 | 33.45 | 10 | YDGSPDGHT | 67.1 |
| S6AR | 7ON8 | 33.47 | 10 | YDGSPDGHT | 75.3 |
| S6AC | 7ONA | 32.87 | 10 | YDGSPDGHT | 87.2 |
| S6BE | 7ONC, 7ONE | 30.46 | 6 | YDGNTH | > 95 |
| S6BR | 7Q58 | 29.67 | 6 | YDGNTH | 67.7 |
| S6BC | 7PVP | 30.76 | 6 | YDGNTH | 92.6 |
| S6BE-L1 | 7ONG | 32.79 | 8 | YDGTGYNT | 60.6 |
| S6BE-L2 | 7ON7 | 68.04 | 11 | YDGSPDYSTGT | 56 |
| S6BE-L3 | 7ONH | 68.97 | 14 | FATLETESGELVE | 51.7 |
| S6AC-L1 | N.A. | 31.94 | 8 | YDGTGYNT | 88.7 |
| S6AC-L2 | N.A. | 33.55 | 11 | YDGSPDYSTGT | 71.6/82.9 |
| S6AC-L3 | N.A. | 33.73 | 14 | FATLETESGELVPE | 66.0/85.8 |
| S6AC-LB | 8PJR | 30.43 | 6 | YDGNTH | > 95 |
| S6BE-3HH | 7OPU, 7OP4, 7OPV | 30.43 | 6 | YDGNTH | > 95 |
| S6BE-6HH | N.A | 29.95 | 6 | YDGNTH | N.A |

The CD signal at 233 nm was followed from 5 to 95 °C with intervals of 0.2 °C. The data was analyzed with a Python script, which fits a Boltzmann-sigmoid equation and extracts the midpoint Tm (Equation 1). The parameters are analogous to those described in a previous publication by Mylemans et al.7 Some data showed a two-state melting curve. In such cases, a two-state version of the above function was used to fit the data. This equation is obtained following the linear addition of a second exponential term (Equation 2).

**Equation 1:**

$$F\left( x \right)=\alpha_{N}+\beta{}_{N}\left( T+x \right)+\left( \alpha_{D}+\beta{}_{N}\left( T+x \right) \right)\frac{exp\left( \frac{-DH_{m}\left( 1-\frac{T+x}{T+T_{m}} \right)}{R\left( T+x \right)} \right)}{1+exp\left( \frac{-DH_{m}\left( 1-\frac{T+x}{T+T_{m}} \right)}{R\left( T+x \right)} \right)}$$

**Equation 2:**

$$F\left( x \right)=\alpha_{N}+\left( \alpha_{D}+\beta_{D}\left( T+x \right) \right)\left( \frac{n_{1}exp\left( \frac{-DH_{m1}\left( 1-\frac{T+z}{T+T_{m1}} \right)}{R\left( T+x \right)} \right)}{1+exp\left( \frac{-DH_{m1}\left( 1-\frac{T+z}{T+T_{m1}} \right)}{R\left( T+x \right)} \right)}+\frac{\left( 1-n_{1} \right)exp\left( \frac{-DH_{m2}\left( 1-\frac{T+z}{T+T_{m2}} \right)}{R\left( T+x \right)} \right)}{1+exp\left( \frac{-DH_{m2}\left( 1-\frac{T+z}{T+T_{m2}} \right)}{R\left( T+x \right)} \right)} \right)$$

**Table S2:** SAKe crystallization conditions (excluding pH-induced crystals).

| Protein | PDB code | Buffer |
| --- | --- | --- |
| S6BE | 7ONC | 0.2 M Potassium formate, 20 %(w/v) PEG 3350 |
| S6BC | 7PVP | 0.2 M Ammonium phosphate, 0.1 M tris pH 8.5, 50% (v/v) MPD |
| S6BR | 7Q58 | 0.09 M HEPES sodium salt pH 7.5, 1.26 M tri-Sodium citrate,  10% (v/v) Glycerol |
| S6AE | 7ON6 | 0.2 M tri-potassium citrate, 1.6 M AmSO4 |
| S6AC | 7ONA | 0.19 M Calcium chloride, 0.095 M HEPES sodium salt pH 7.5,  26.6% (v/v) PEG 400, 5% (v/v) Glycerol |
| S6AR | 7ON8 | 58% (v/v) MPD, 20% (v/v) Glycerol, 0.085 M HEPES pH 7.5 |
| S6BE-L1 | 7ONG | 0.2 M Calcium acetate pH 6.5, 0.1 M Sodium cacodylate,  40% (v/v) PEG 300 |
| S6BE-L2 | 7ON7 | 0.2 M Magnesium chloride, 0.1 M Tris pH 8.5,  30% (w/v) PEG 4000 |
| S6BE-L3 | 7ONH | 0.1 M Tris pH 8.0, 2.4 M Ammonium sulfate |
| S6BE-3HH | 7OP4 | 0.1 M MES, 0.9 M Na phosphate, 0.9 M K phosphate, pH 6.5 |
| S6BE-3HH (alt) | 7OPV | 0.1 M MES, 1.2 M tri-Na citrate, 0.995 mM Cu(NO3)2, pH 6.5 |
| S6AC-LB | 8PJR | 0.2 M Magnesium formate pH 5.9, 20% PEG 3350 |

**Table S3:** Crystallographic data table for A-type SAKe.

| Parameters and statistics | S6AE (7ON6) | S6AR (7ON8) | S6AC (7ONA) |
| --- | --- | --- | --- |
| Data collection | | | |
| Space group | P 31 2 1 | P 21 21 21 | C 1 2 1 |
| Unit cell, a, b, c (Å) | 82.767 82.767 | 45.986 63.249 | 88.3804 74.8725 46.938 |
| α, β, γ (°) | 90.023 90 90 120 | 99.567 90 90 90 | 90 122.079 90 |
| Resolution range (Å) | 38.12 - 1.35 (1.398 - 1.35) | 39.12 - 1.5 (1.554 - 1.5) | 39.77 - 1.45 (1.502 - 1.45) |
| Total reflections | 1541949 (142973) | 618399 (60166) | 307460 (28170) |
| Unique reflections | 78555 (7776) | 47304 (4657) | 45754 (4441) |
| Multiplicity | 19.6 (18.4) | 13.1 (12.9) | 6.7 (6.3) |
| Completeness (%) | 99.99 (100.00) | 99.98 (100.00) | 99.37 (98.36) |
| I/σI | 50.48 (6.70) | 26.77 (2.58) | 24.69 (3.07) |
| Rmerge | 0.03554 (0.5203) | 0.06382 (1.053) | 0.03998 (0.5624) |
| CC1/2 | 1 (0.966) | 1 (0.818) | 1 (0.899) |
| Refinement | | | |
| Rwork/Rfree (%) | 13.7 / 15.8 | 17.6 / 18.2 | 17.0 / 18.5 |
| Reflections used in refinement | 78552 (7776) | 47302 (4657) | 45535 (4436) |
| R.m.s deviations: |  |  |  |
| bond lengths (Å) | 0.009 | 0.011 | 0.008 |
| bond angles (°) | 1.1 | 1.22 | 1.03 |
| Ramachandran analysis: |  |  |  |
| Favoured (%) | 97.36 | 95.3 | 97.72 |
| Allowed (%) | 2.64 | 4.7 | 2.28 |
| Outliers (%) | 0 | 0 | 0 |
| Number of atoms | 2715 | 2477 | 2363 |
| Protein (average B-factor, Å²) | 17.7 | 17.76 | 19.03 |
| Solvent (average B-factor, Å²) | 33.49 | 29.84 | 28.09 |
| Ligands (average B-factor, Å²) | 29.27 | N.A. | 20.96 |
| Mean/Wilson B-factor (Å²) | 19.94 / 14.63 | 18.82 / 15.39 | 19.92 / 16.09 |
| Number of TLS groups | N.A. | 6 | N.A. |

**Table S4:** Crystallographic data table for B-type SAKe.

| Parameters and statistics | S6BE (normal) (7ONC) | S6BR (7Q58) | S6BC (7PVP) | S6BE (self assembled) (7ONE) |
| --- | --- | --- | --- | --- |
| Data collection |  |  |  |  |
| Space group | P 1 | P 1 21 1 | P 1 | P 1 |
| Unit cell, a, b, c (Å) | 34.981 46.779 46.809 | 34.384 50.322 71.324 | 34.342 46.858 46.9107 | 35.0258 46.5119 46.5057 |
| α, β, γ (°) | 119.972 90.014 90.023 | 90 90 90 | 60.0574 89.9891 89.966 | 60.0511 89.988 89.9399 |
| Resolution range (Å) | 40.55 - 1.49 (1.543 - 1.49) | 28.39 - 1.3 (1.346 - 1.3) | 40.65 - 1.8 (1.864 - 1.8) | 40.3 - 1.3 (1.346 - 1.3) |
| Total reflections | 41309 (14341) | 396946 (37474) | 77256 (7431) | 191819 (16769) |
| Unique reflections | 40395 (3985) | 59917 (5969) | 22639 (2225) | 57085 (6238) |
| Multiplicity | 3.5 (3.6) | 6.6 (6.3) | 3.4 (3.3) | 3.4 (3.0) |
| Completeness (%) | 96.08 (95.04) | 99.90 (99.85) | 96.83 (95.68) | 99.68 (99.97) |
| I/σI | 52.46 (37.04) | 23.57 (3.33) | 10.48 (2.62) | 44.08 (22.29) |
| Rmerge | 0.02053 (0.02645) | 0.0405 (0.5952) | 0.07549 (0.4295) | 0.01945 (0.04152) |
| CC1/2 | 0.999 (0.999) | 1 (0.902) | 0.996 (0.889) | 0.999 (0.996) |
| Refinement |  |  |  |  |
| Rwork/Rfree (%) | 15.8 / 18.4 | 14.9 / 17.6 | 17.3 / 21.4 | 13.6 / 16.3 |
| Reflections used in refinement | 40387 (3985) | 59909 (5962) | 22711 (2216) | 62372 (6238) |
| R.m.s deviations: |  |  |  |  |
| bond lengths (Å) | 0.012 | 0.009 | 0.015 | 0.009 |
| bond angles (°) | 1.11 | 1.42 | 1.17 | 1.06 |
| Ramachandran analysis: |  |  |  |  |
| Favoured (%) | 97.85 | 98.92 | 97.85 | 95.34 |
| Allowed (%) | 1.03 | 1.08 | 1.17 | 4.66 |
| Outliers (%) | 0 | 0 | 0 | 0 |
| Number of atoms | 2341 | 2344 | 2201 | 2401 |
| Protein (average B-factor, Å²) | 12.99 | 13.85 | 17.63 | 17.96 |
| Solvent (average B-factor, Å²) | 25.25 | 25.9 | 29.56 | 27.38 |
| Ligands (average B-factor, Å²) | N.A. | N.A. | N.A. | N.A. |
| Mean/Wilson B-factor (Å²) | 14.36 / 11.37 | 15.39 / 13.69 | 18.83 / 19.83 | 15.39 / 11.68 |
| Number of TLS groups | 9 | N.A. | 12 | N.A. |

**Table S5:** Crystallographic data table for SAKe loop variants.

| Parameters and statistics | S6BE-L1 (7ONG) | S6BEL2 (7ON7) | S6BE-L3 (7ONH) | S6AC-LB (8PJR) |
| --- | --- | --- | --- | --- |
| Data collection |  |  |  |  |
| Space group | C 1 2 1 | P 1 21 1 | P 1 1 | P 1 1 |
| Unit cell, a, b, c (Å) | 43.7842 75.7993 72.228 | 79.9579 45.4007 | 90.624 90.624 90.663 | 34.872 47.703 74.797 |
| α, β, γ (°) | 90 90.003 90 | 81.6208 90 114.721 90 | 90 90 90 | 90 90.12 90 |
| Resolution range (Å) | 37.9 - 1.95 (2.02 - 1.95) | 39.86 - 1.95 (2.02 - 1.95) | 40.53 - 1.65 (1.709 - 1.65) | 25.48 - 1.71 (1.771 - 1.71) |
| Total reflections | 119377 (11749) | 247539 (17550) | 1181876 (80392) | 182965 (18513) |
| Unique reflections | 16946 (1716) | 38757 (3819) | 87019 (8786) | 26294 (2604) |
| Multiplicity | 7.0 (7.1) | 6.4 (6.5) | 13.6 (10.2) | 7.0 (7.1) |
| Completeness (%) | 99.89 (99.94) | 99.4 (98.30) | 99.96 (99.91) | 98.01 (97.67) |
| I/σI | 10.19 (2.75) | 6.6 (2.1) | 48.64 (12.94) | 21.69 (10.55) |
| Rmerge | 0.1459 (0.7654) | 0.135 (0.675) | 0.03584 (0.1437) | 0.07843 (0.4605) |
| CC1/2 | 0.996 (0.868) | 0.993 (0.943) | 1 (0.991) | 0.996 (0.958) |
| Refinement |  |  |  |  |
| Rwork/Rfree (%) | 19.2 / 21.8 | 22.8 / 27.9 | 16.0 / 17.7 | 19.4 / 20.4 |
| Reflections used in refinement | 17244 (1715) | 38757 (3819) | 87937 (8786) | 26243 (2603) |
| R.m.s deviations: |  |  |  |  |
| bond lengths (Å) | 0.002 | 0.003 | 0.004 | 0.008 |
| bond angles (°) | 0.53 | 0.54 | 0.73 | 1.56 |
| Ramachandran analysis: |  |  |  |  |
| Favoured (%) | 95.29 | 97.68 | 98.09 | 96.06 |
| Allowed (%) | 4.71 | 2.32 | 1.91 | 3.94 |
| Outliers (%) | 0 | 0 | 0 | 0 |
| Number of atoms | 2305 | 4660 | 5506 | 2237 |
| Protein (average B-factor, Å²) | 20.45 | 32.73 | 14.17 | 15.45 |
| Solvent (average B-factor, Å²) | 23.15 | 37.43 | 39.05 | 27.18 |
| Ligands (average B-factor, Å²) | 26.2 | N.A. | 27.2 | N.A. |
| Mean/Wilson B-factor (Å²) | 20.63 / 19.37 | 33.32 / 22.52 | 16.3 / 12.93 | 13.6 / 13.7 |
| Number of TLS groups | 1 | 2 | 13 | 8 |

**Table S6:** Crystallographic data table for S6BE-3HH SAKe.

| Parameters and statistics | S6BE-3HH (7OPV) | S6BE-3HH (self-assembled) (7OPU) | S6BE-3HH (alternative packing) (7OPV) |
| --- | --- | --- | --- |
| Data collection |  |  |  |
| Space group | C 1 2 1 | C 1 2 1 | P 1 1 |
| Unit cell, a, b, c (Å) | 44.049 76.29 67.987 | 76.4486 44.1528 68.326 | 46.9614 47.1524 70.876 |
| α, β, γ (°) | 90 90.021 90 | 90 90.0535 90 | 72.1828 76.4037 60.1756 |
| Resolution range (Å) | 38.15 - 1.52 (1.574 - 1.52) | 68.33 - 1.7 (1.761 - 1.7) | 40.54 - 1.95 (2.02 - 1.95) |
| Total reflections | 234771 (23764) | 169862 (16090) | 120140 (11837) |
| Unique reflections | 33235 (3298) | 25255 (2491) | 34968 (3389) |
| Multiplicity | 7.1 (7.2) | 6.7 (6.5) | 3.4 (3.5) |
| Completeness (%) | 96.01 (95.04) | 99.79 (99.52) | 96.00 (94.43) |
| I/σI | 30.43 (8.87) | 35.05 (14.57) | 35.05 (14.57) |
| Rmerge | 0.03691 (0.207) | 0.0454 (0.5517) | 0.03621 (0.06553) |
| CC1/2 | 0.999 (0.99) | 1 (0.878) | 0.995 (0.29) |
| Refinement |  |  |  |
| Rwork/Rfree (%) | 18.6 / 20.2 | 18.4 / 22.6 | 15.9 / 20.4 |
| Reflections used in refinement | 33207 (3293) | 25216 (2479) | 34944 (3388) |
| R.m.s deviations: |  |  |  |
| bond lengths (Å) | 0.006 | 0.006 | 0.007 |
| bond angles (°) | 0.93 | 0.85 | 0.79 |
| Ramachandran analysis: |  |  |  |
| Favoured (%) | 97.12 | 97.49 | 96.24 |
| Allowed (%) | 2.88 | 2.51 | 3.76 |
| Outliers (%) | 0 | 0 | 0 |
| Number of atoms | 2320 | 2268 | 4547 |
| Protein (average B-factor, Å²) | 19.64 | 25.86 | 20.59 |
| Solvent (average B-factor, Å²) | 30.6 | 35.06 | 32.25 |
| Ligands (average B-factor, Å²) | N.A. | N.A. | N.A. |
| Mean/Wilson B-factor (Å²) | 20.72 / 16.85 | 24.65 / 23.2 | 21.71 / 19.29 |
| Number of TLS groups | 5 | 6 | 18 |

**III. SUPPLEMENTARY SCRIPTS**

**Random loop generation script (Part 1)**

import sys

\# Calculate the fraction of occurrence of each amino acid in the iCAN database TXT file.

args = sys.argv

file = args[1]

with open(file) as f:

text = f.read()

def count_char(text, char):

count = 0

for c in text:

if c == char:

count += 1

return count

for char in "ACDEFGHIKLMNPQRSTVWY":

perc = 100 * count_char(text, char) / len(text)

print("{0} - {1}%".format(char, round(perc, 2)))

**Random loop generation script (Part 2)**

\# Generate a random string of specific characters

from random import random

from bisect import bisect

import string

\# Weights for each amino acid in a cumulative distribution:

#random.random() to pick a random float 0.0 <= x < total

#search the distribution with bisect.bisect:

#Randomly generate a number between 0 and 1:

b = int(input('Amount: '))

l = int(input('Length: '))

def weighted_choice(choices):

values, weights = zip(*choices)

total = 0

cum_weights = []

for w in weights:

total += w

cum_weights.append(total)

x = random() * total

i = bisect(cum_weights, x)

return values[i]

def weighted_sequence(length=l):

return ''.join((weighted_choice([("A",8.04), ("C",0.58),

("D",7.85), ("E",1.9), ("F",1.7), ("G",12.86), ("H",1.15),

("I",4.81), ("K",3.43), ("L",2.77), ("M",2.71), ("N",3.87),

("P",2.32), ("Q",0.71), ("R",5.19), ("S",13.13), ("T",7.51),

("V",5.65), ("W",2.16), ("Y",11.63),])

for i in range(length)))

c = 0

while c < b:

sequence = weighted_sequence()

\# print(sequence)

if "Y" in sequence and "H" in sequence:

print("Weighted sequence ", sequence)

c +=
